# DNA Modifications Enabling Proximity Biotinylation

**DOI:** 10.1101/2022.10.02.510503

**Authors:** Brandon Wilbanks, Keenan Pearson, Shane R. Byrne, Laura B. Bickart, Peter C. Dedon, L. James Maher

## Abstract

Advances in peroxidase- and biotin ligase-mediated signal amplification have enabled high-resolution subcellular mapping of endogenous RNA localization and protein-protein interactions. Application of these technologies has been limited to RNA and proteins because of the reactive groups required for biotinylation in each context. Here we report several novel methods for proximity biotinylation of exogenous oligodeoxyribonucleotides by application of well-established and convenient enzymatic tools. We describe approaches using simple and efficient conjugation chemistries to modify deoxyribonucleotides with “antennae” sensitive to phenoxy radical or biotinoyl-5’-adenylate. In addition, we report chemical details of a previously undescribed adduct between tryptophan and a phenoxy radical group. These developments have potential application in the selection of exogenous nucleic acids capable of unaided entry into living cells.

---

With the development of the biotin ligase BioID in 2012 [1], proximity biotinylation has become a powerful tool for the mapping of protein-protein interactions and protein localization in living cells. More recent advances have enabled the mapping of protein and RNA localization across entire subcellular compartments with the ascorbate peroxidase APEX or its optimized form, APEX2 [2-4]. Despite the widespread success of these approaches, an equivalent method for mapping DNA localization and DNA-protein interactions has yet to be achieved. BioID and TurboID Biotin ligases catalyze synthesis of the highly reactive biotinoyl-5’-AMP intermediate from biotin and ATP, allowing proximity biotinylation of primary amine nucleophiles as reactive targets [1,5]. While such targets exist in proteins on lysine residues and N-termini, none are found in nucleic acids. APEX and APEX2, on the other hand, catalyze production of short-lived reactive phenoxy radical intermediates from biotin tyramide (BT) in the presence of H_2_O_2_. Though proximity biotinylation by peroxidases has the impressive advantage of dual protein and RNA radical biotinylation, this approach cannot be used directly for biotinylation of unmodified DNA [6].

We have previously shown that peroxidase-based biotinylation of oligodeoxyribonucleotides with BT is enabled by DNA conjugation with 5’ fluorescein and we described structural details of BT-fluorescein adducts [6]. We demonstrated that an in vitro streptavidin gel shift assay is capable of detecting biotinylation of both RNA and 5’-fluorescein-modified DNA oligonucleotides and, importantly, provided molecular insights into differences in RNA and DNA biotinylation. Here, we expand the toolbox of DNA proximity biotinylation methods with new approaches suitable for peroxidases and, for the first time, biotin ligases. We envision application of these approaches for selection of exogenous DNAs capable of unaided entry into living cells.

It is known that tyrosine residues are protein reactive sites biotinylated by APEX2-generated BT phenoxy radicals [7]. To determine if other biological functional groups are reactive with BT phenoxy radicals, we devised a competition assay for the assessment of all other natural amino acids. In this assay, a streptavidin gel shift is used to detect horseradish peroxidase (HRP)and H_2_O_2_-dependent radical biotinylation of 5’-fluorescein modified oligodeoxyribonucleotide LJM-6132 [6]. Increasing concentrations of individual competing amino acids are added to the reaction. Amino acid reaction with BT radicals is detected by quenching of LJM-6132 biotinylation. This results in a quantifiable inhibition of LJM-6132 gel shifts.

We first observed that excess fluorescein reduces biotinylation-dependent streptavidin gel shifts as expected, validating that this assay is sensitive to competition for radicals (Fig. 1). We then tested all 20 natural amino acids in competition with fluorescein-modified LJM-6132 at 100-, 500-, and 1000fold excess to LJM-6132 (Fig. S1-S3). As expected, excess tyrosine competes for phenoxy radicals and reduces the gel shift observed under competitor-free conditions (Fig. 1). Notably, our investigation revealed that tryptophan also quenches the reaction. Tryptophan reactivity with phenoxy radicals has not been previously described. The remaining 18 amino acids, exemplified here by glycine, were unreactive (Fig. 1; Figs. S1-S3). These results suggest that both phenol and indole functional groups might serve as simple biotin tyramide-receptive “antennae” for proximity biotinylation of appropriately conjugated DNA oligonucleotides.

**Figure 1.**
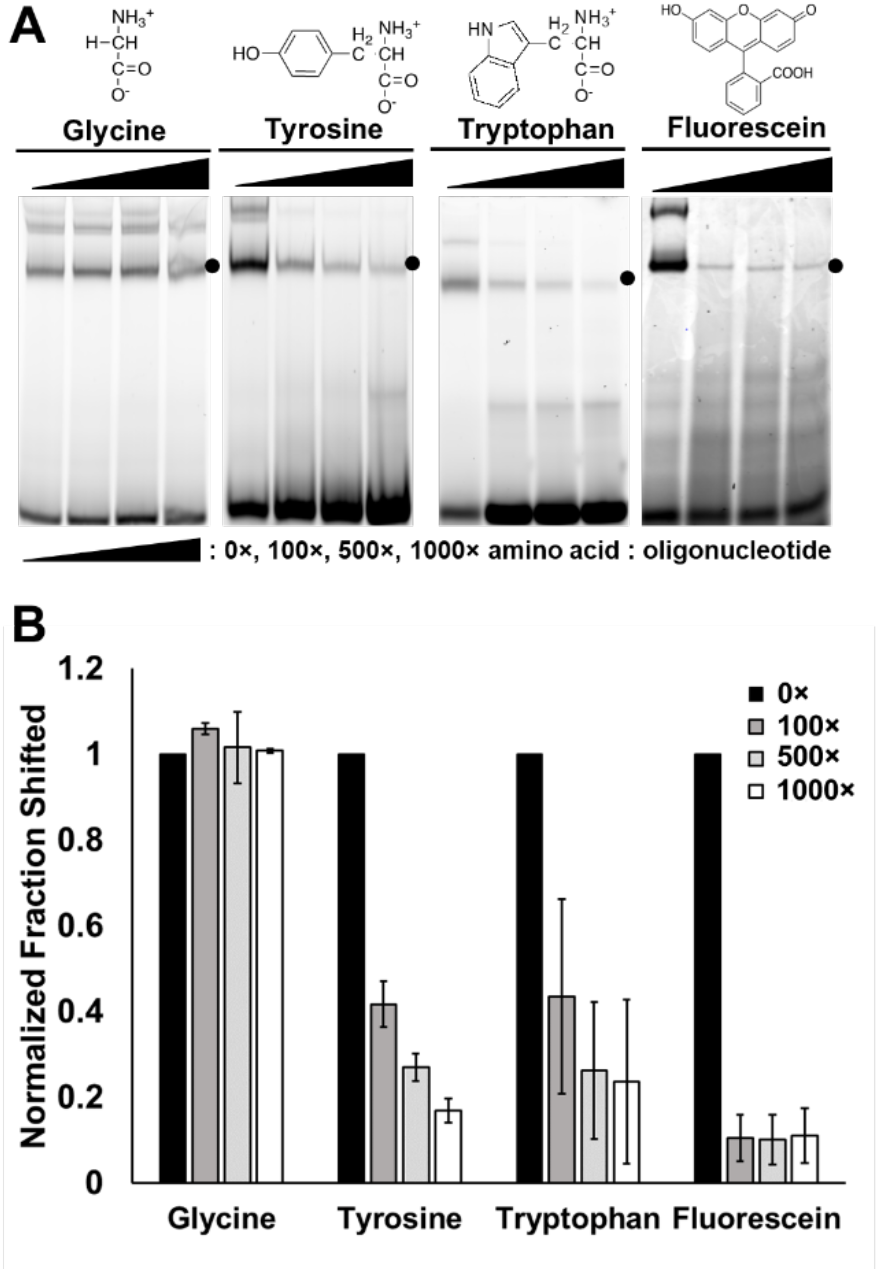
Amino acids tyrosine and tryptophan compete for phenoxy radical biotin tyramide intermediates to quench biotinylation. A) Excess amino acids or fluorescein are added to *in vitro* HRP-mediated biotinylation reaction to monitor competition for radical reactivity. Covalently biotinylated oligonucleotides are detected by streptavidin gel shift, which is quenched in the presence of reactive competitors. B) Quantification of gel shifted material in the presence of various amino acids or fluorescein.

Having screened all natural amino acids for phenoxy radical reactivity, we set out test tyrosine and tryptophan as DNA conjugates in place of fluorescein in our established gel shift assay of phenoxy radical biotinylation. Tyrosine hydrazide was conjugated to 5’ aromatic aldehyde-modified oligodeoxyribonucleotide LJM-6542 to generate LJM-6542-5’Tyr (Fig. 2A). Tyrosine-conjugated oligonucleotides were purified from denaturing polyacrylamide gels (Fig. S4A). Unexpectedly, we first found that the 5’ aromatic aldehyde modification of LJM6542 was itself sensitive to phenoxy BT radicals, though only approximately 15-20% as reactive as 5’ fluorescein (Fig. 2B, lane 7; Fig. S4B, lane 7; Fig. S4C). The LJM-6542-5’Tyr conjugate was also slightly less reactive than the 5’ fluorescein of LJM-6132, though significantly more reactive than LJM-6542 (Fig. 2C, lane 7; S4B-C). Biotinylation of both the 5’ aromatic aldehyde of LJM-6542 and LJM-6542-5’Tyr requires the simultaneous presence of BT, H_2_O_2_, and HRP, confirming that the catalyzed reaction is similar to that for peroxidase proximity labeling of proteins [7]. Thus, despite being approximately 25% less reactive than 5’ fluorescein, 5’ tyrosine modifications present a useful alternative for DNA oligonucleotide biotinylation in cases where use of a fluorescent substrate is incompatible with the required application. This may be helpful in fluorescence microscopy applications when other fluorescein-labeled reagents are required.

**Figure 2.**
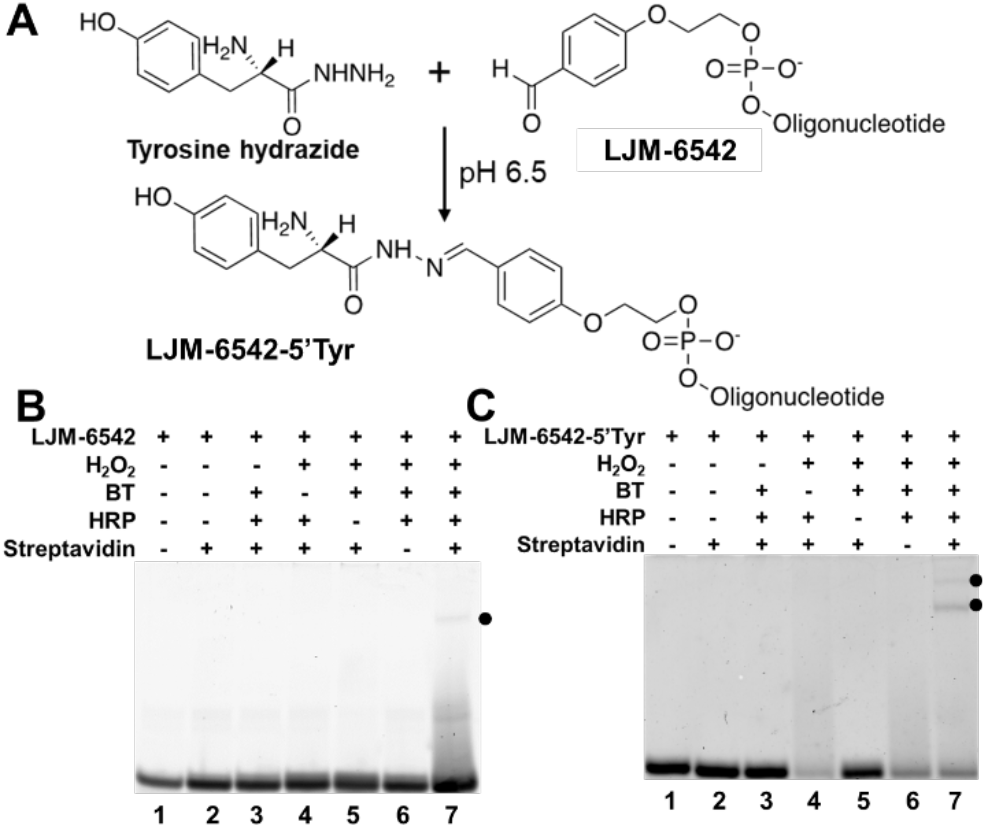
Tyrosine DNA oligonucleotide conjugates react with phenoxy biotin tyramide radicals. A) Reaction scheme for covalent conjugation of tyrosine hydrazide and a 5’ aromatic aldehyde-modified oligonucleotide. B) Biotinylation and streptavidin gel shift of 5’ aldehyde modified oligonucleotide LJM-6542 (black dot) occurs only in the presence of H2O2, HRP, and BT. C) Biotinylation and streptavidin gel shift of 5’ tyrosine-conjugated oligonucleotide LJM-6542 (black dot) occurs only in the presence of H2O2, HRP, and BT. DNA is visualized by SYBR Gold.

We next tested a 5’ tryptophan conjugate as an alternative to fluorescein to facilitate DNA oligonucleotide biotinylation. A N*α*-Boc-l-tryptophan N-hydroxysuccinimide (NHS) ester (Trp-NHS) was coupled to the 5’ primary amine modification of oligodeoxyribonucleotide LJM-6543 (Fig. 3A). The conjugated product, LJM-6543-5’Trp, was purified by denaturing polyacrylamide gel electrophoresis (Fig. S5A). 5’ conjugation of Trp-NHS was further confirmed by high resolution LC-Orbitrap-MS to identify the predicted product (Fig. S5B, D). LJM-6543-5’Trp was amenable to phenoxy radical biotinylation and streptavidin gel shift indicated approximately 50% reactivity compared to the fluorescein conjugate (Fig. 3B, lane 7; Fig. 3C). Remarkably, we found that biotinylation of LJM6543-5’Trp occurred even in the absence of HRP. requiring only BT and H_2_O_2_ (Fig. 3B, lane 6; Fig. 3C). This reaction, enhanced by the addition of peroxidase, could serve as a tool for non-enzymatic biotinylation of Trp-containing DNA conjugates. These results also suggest that proximity biotinylation by APEX2 might be subject to non-enzymatic background biotinylation upon prolonged cell incubation with biotin tyramide and H_2_O_2_.

**Figure 3.**
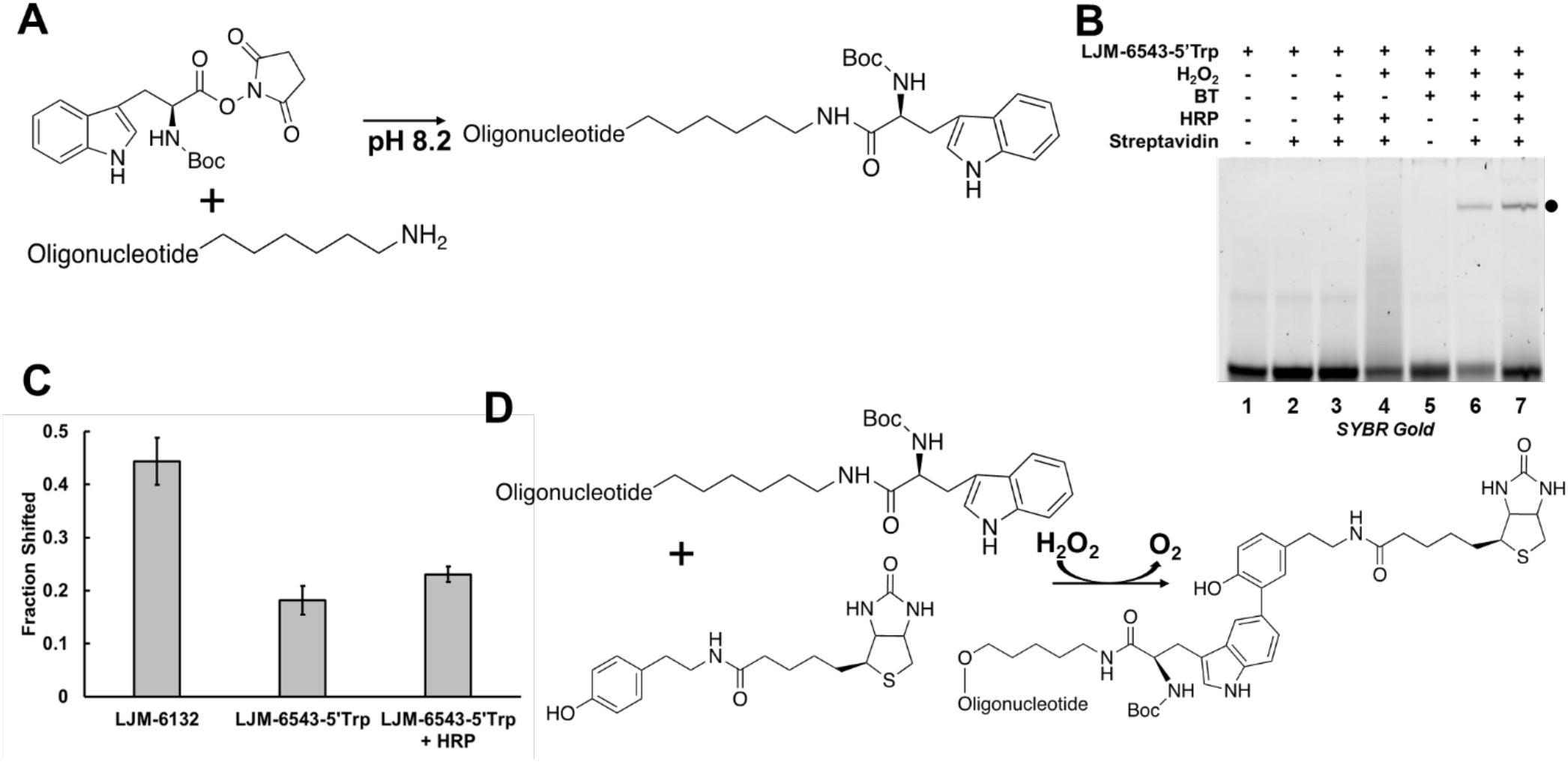
Tryptophan-DNA oligonucleotide conjugates react with phenoxy biotin tyramide radicals. A) Reaction scheme for conjugation of tryptophan NHS ester to 5’ primary amine-modified oligonucleotides. B) Phenoxy radical biotinylation of LJM-6543-5’Trp occurs in the presence of H2O2 and BT alone (lane 6) but is enhanced by the addition of HRP (lane 7). C) Quantification of gel shifts of LJM-6542-5’Trp in with or without acceleration by HRP versus 5’ fluorescein modified LJM-6132. D) Proposed reaction scheme of unexpected HRP-free biotinylation of 5’ tryptophan-modified oligonucleotides.

High resolution LC-Orbitrap-MS analysis of the HRP-free biotinylation product further confirmed the unexpected uncatalyzed biotinylation of LJM-6543-5’Trp. We identified a previously unreported 5’Trp-BT adduct (Fig. S5C, 5E) and propose a novel reaction between tryptophan and BT that can be accelerated by peroxidase (Fig. 3D). This reaction can occur on 5’-Trp conjugated oligonucleotides or with free tryptophan, as evidenced by the ability of free tryptophan to quench biotinylation in a competition assay (Fig. S6A). The reaction therefore does not depend on the N*α*-Boc group present in the 5’Trp conjugate that we generated here. Tryptophan biotinylation does, however, require the tyramide group of BT as shown by the absence of a gel shifted species if the reaction is carried out with biotin rather than BT (Fig. S6B).

We next studied the possibility of expanding the suite of DNA proximity biotinylation tools from those compatible only with peroxidases to approaches suitable for biotin ligases. Proximity biotinylation by stably-expressed biotin ligases relies only on exogenous addition of biotin to cells, unlike biotinylation by APEX2, which requires the addition of toxic H_2_O_2_. In the presence of endogenous ATP, biotin ligases generate highly reactive biotin-5’-adenylate intermediates that then react with proximal nucleophiles such as primary amines. Recently, directed evolution was used to develop the highly promiscuous biotin ligase TurboID which significantly outperforms previously established enzymes such as BioID [8]. We therefore expressed and purified TurboID for *in vitro* reactions to model oligonucleotide biotinylation by biotin ligase (Fig. S7). To first ensure that TurboID was enzymatically active and capable of labeling proteins, we combined TurboID, ATP, and biotin *in vitro* with HRP now serving as a convenient biotinylation target. Western blot analysis of the products using AlexaFluor647-streptavidin to mark labeled proteins revealed both promiscuous intramolecular biotinylation of TurboID and intermolecular biotinylation of HRP, confirming TurboID activity (Fig. S8).

The parent enzyme of TurboID, BirA, transfers biotin to a target lysine through a reactive biotinyl-5’-adenylate intermediate [9,10]. Because DNA lacks aliphatic primary amines, we reasoned that the addition of convenient and commercially available amino modifications would allow oligonucleotide biotinylation by TurboID (Fig. 4A). We first demonstrated this using streptavidin gel shift assays to detect biotinylation of a 5’C_6_-amino modified oligonucleotide in the presence of TurboID, ATP, and biotin (Fig. S9A-B). Oligonucleotides lacking the primary amine modification are not detectably biotinylated (Fig. 4B, lane 1). Multiple amino-containing oligonucleotide modifications are commercially available. We compared biotinylation of oligonucleotides containing primary amines on spacers (C_6_ and C_12_) and those with internal amino-modified deoxythymidine bases (Fig. 4B, lanes 2-4, Fig. S9C-D). All variants carrying primary amines were similarly biotinylated (Fig. S9E). To further demonstrate biotinyl-5’-adenylate specificity for primary amines, a competition assay was performed by adding excess glycine to the reaction mixture. Competition between amino-modified DNA and glycine for reactive intermediates resulted in a diminished gel shift (Fig. S10A). We also demonstrated that observed gel shifts are dependent on the predicted streptavidin-biotin interaction by blocking streptavidin with excess biotin prior to adding the oligonucleotide, resulting in a reduced gel shifts (Fig. S10B).

**Figure 4.**
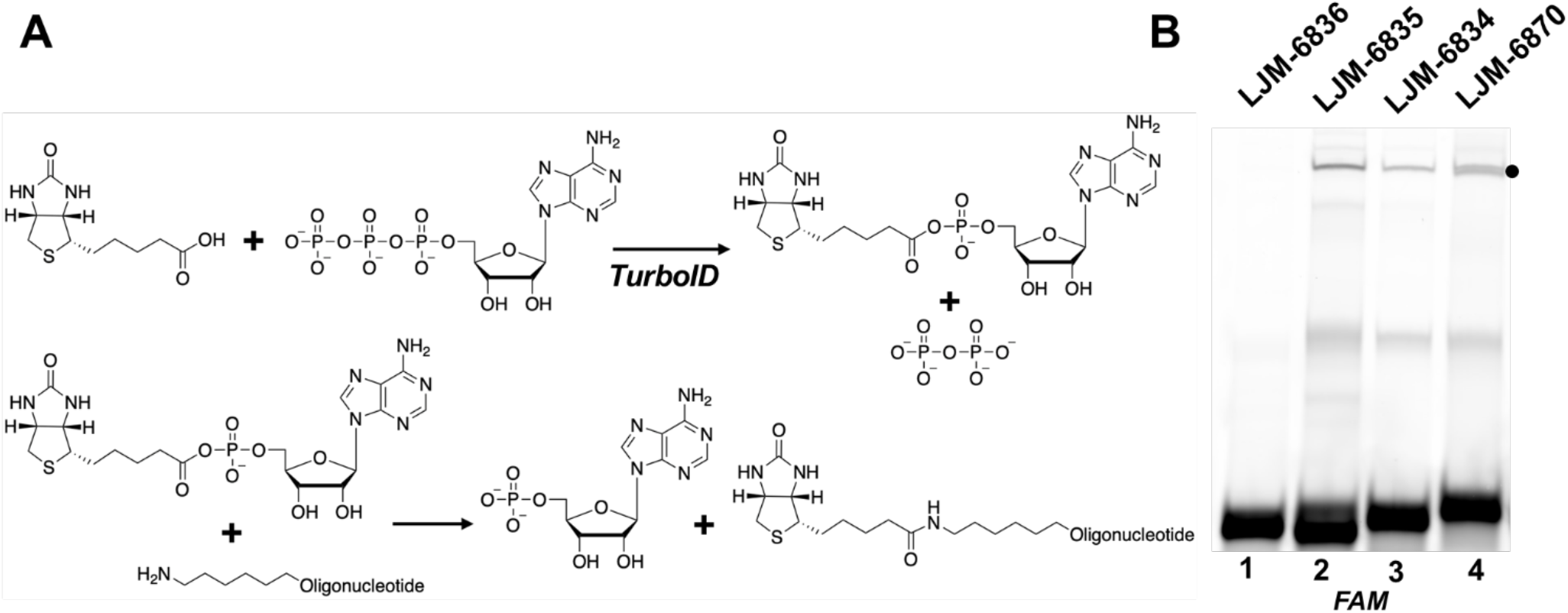
DNA carrying a primary amine is a substrate for biotinylation by biotin ligase. A) Schematic of reaction between ATP and biotin catalyzed by TurboID yielding biotinyl-5’-adenylate, which subsequently reacts with a primary amine-modified oligonucleotide. B) Comparison of streptavidin gel shifts (black dot) for an oligonucleotide containing no primary amine (LJM-6836), an oligonucleotide containing a 5’ primary amine with a C12 spacer (LJM-6835), an oligonucleotide containing a 5’ primary amine with a C6 spacer (LJM-6834), and an oligonucleotide containing a 5’ primary amine with a C6 spacer and two internal primary amine modified deoxythymidine nucleotides (LJM-6870).

As has been discussed for other S_N_2 reactions involving biological nucleophiles [11], reaction products may have different stabilities that obscure their formation. The DNA molecules in our studies possess one or more primary alcohol residues that could act as nucleophiles to produce esters upon reaction with biotinyl-5’-adenylate. Such esters may be unstable under these conditions, explaining the absence of detectable biotinylation of unmodified DNA. The aromatic amino groups of DNA bases are insufficiently nucleophilic to react [12].

We have demonstrated several novel methods for biotinylation of exogenous DNAs. Methods for peroxidase-based biotinylation confirm our previous observation that fluorescein is readily biotinylated by BT. We now show the utility of tyrosine-DNA conjugates for biotinylation using BT. Tryptophan could also be biotinylated by BT in a peroxidase-catalyzed reaction. Remarkably, tryptophan was also detectably biotinylated by BT in the absence of peroxidase. Therefore, tryptophan may be less desirable as a conjugate “antenna” due to the possibility of background biotinylation unrelated to peroxidase activity. We also show that biotin ligase-dependent biotinylation can be extended to amino-modified exogenous DNAs in the presence of biotin and ATP. The addition of biotin ligase-based tools allows proximity biotinylation of nucleic acids without the need for toxic H_2_O_2_. We envision application of these methods for selection of exogenous DNA oligonucleotides capable of homing to desired subcellular compartments.

## ASSOCIATED CONTENT

### Supporting Information

Supporting information includes materials and methods as well as supplemental figures.

### Author Contributions

Project conception: B.W., L.J.M., P.D.

Manuscript preparation: B.W., K.P., and L.J.M.

Experimental work: B.W., K.P., S.B., and L.B.B.

### Funding Sources

This work was supported by NIH grant GM128579 (L.J.M.), the Mayo Clinic Graduate School of Biomedical Sciences (B.W., K.P.), and an NSF graduate fellowship (B.W.).

## ACKNOWLEDGMENT

We thank Alice Ting for the gift of a DNA construct for TURBO biotin ligase expression. We thank members of the Maher laboratory for their assistance with their work.

## TOC FIGURE

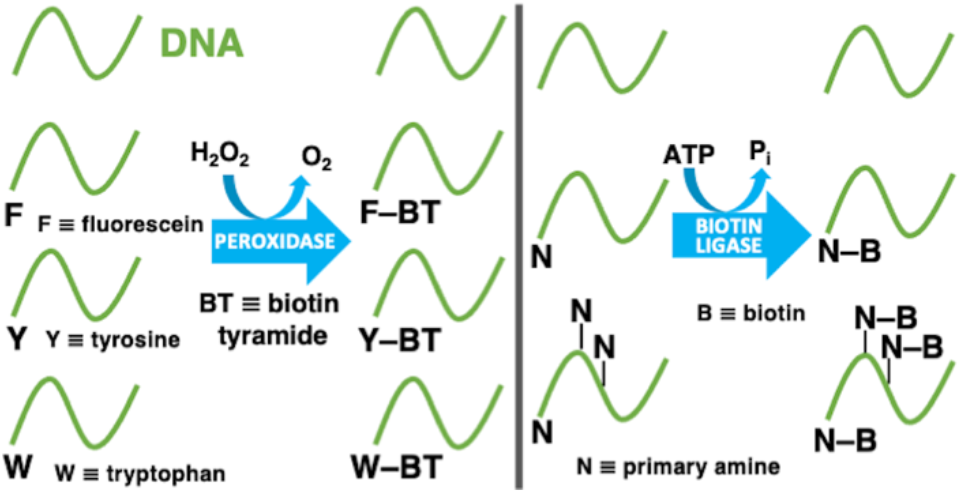

