## Supplemental for "DNA Modifications Enabling Proximity Biotinylation"

**Table of Contents**

**1.0 Methods**

1.1 *In vitro* radical biotinylation assay

1.2 Conjugation of 5'-aldehyde modified oligodeoxyribonucleotide with tyrosine hydrazide

1.3 Conjugation of 5'-amine modified oligodeoxyribonucleotide with tryptophan NHS ester

1.4 Mass spectrometry

1.5 Bacterial expression and purification of TurboID protein

1.6 TurboID biotinylation activity assay by Western blot

1.7 TurboID biotinylation assay

**2.0 Supplemental data**

### 1.0 Methods

#### 1.1 *In vitro* radical biotinylation assay

Oligodeoxyribonucleotides were synthesized by Integrated DNA Technologies (Coralville, IA). Radical biotinylation reactions (40 µL) contained the indicated oligonucleotide (2.5 µM), 10 mM potassium phosphate, pH 6, horseradish peroxidase (HRP, 12.5 µM, Thermo Scientific # 31490), freshly prepared $H_2O_2$ (500 µM, Sigma # 216763), and biotin tyramide (100 µM, ApexBio # A8011). When applicable, competitors were included at the indicated concentration and water was subtracted to maintain a constant final reaction volume. Reactions were incubated at 45 ºC for 1.5 h with agitation at 1000 rpm (Benchmark Scientific, model H5000-HC). Reactions were then diluted by addition of 40 µL water and combined with 2 µL proteinase K solution and 18 µL buffer ATL (Qiagen DNEasy Blood and Tissue Extraction kit # 69504). These reactions were incubated at 56 ºC for 1 h with agitation at 1000 rpm. Reactions were then extracted with phenol:chloroform:isoamyl alcohol (25:24:1, 80 µL, Invitrogen # 15593031) and aqueous phases combined with 0.1 volume 3 M sodium acetate, 1 µL glycogen (Thermo Scientific # R0561) and 2.5 volumes ice-cold 100% ethanol. This mixture was chilled on dry ice for 15 min and subjected to centrifugation for 20 min at 16,000 × g. Samples were washed with 150 µL ice-cold 70% ethanol followed by centrifugation for 10 min at 16,000 × g and air drying. Pellets were resuspended in water (15 µL), and either streptavidin (5 µL of 182 µM stock, Invitrogen # 434301) or water (5 µL) was added with incubation for 30 min at 37 ºC with agitation at 1000 rpm. Samples were then analyzed by electrophoresis. For analysis under native conditions, samples were supplemented with loading buffer (Invitrogen # 10482035) and subjected to electrophoresis at 22.5 V/cm for 30 min through native 10% polyacrylamide (19:1 acrylamide:bisacrylamide) gels. For analysis under denaturing conditions, samples were supplemented with formamide (20 µL, Sigma #F9037), heated to 70˚C for 3 min, and cooled on ice before electrophoresis through 10% polyacrylamide (19:1 acrylamide:bisacrylamide,) gels containing 7.5 M urea. Gels were imaged on a Typhoon5 fluorometric imager (Amersham) using the FAM channel.

#### 1.2 Conjugation of 5'-aldehyde-modified oligodeoxyribonucleotide with tyrosine hydrazide

Tyrosine hydrazide (Sigma-Aldrich # 138045) stock was prepared in anhydrous DMSO at 25 mM. 5 µL of tyrosine hydrazide stock was combined with 5 µL of 1 mM LJM-6542, 2.5 µL 0.1 M NaOAc, pH 5.5, and 32.5 µL water. The solution was mixed thoroughly by pipetting and incubated overnight at room temperature.

The reaction was then combined with 1 µL glycogen and 250 µL EtOH, mixed, and chilled on dry ice for 10 min. This was followed by centrifugation at 17,000 × g for 20 min and the supernatant was decanted. The pellet was washed with 500 µL 70% EtOH and centrifuged again for 5 min. Supernatant was again removed and the pellet was air-dried at 37 °C. After drying, the conjugated oligonucleotide was resuspended in 40 µL water.

The precipitated oligonucleotide was combined with 40 µL formamide and heated at 90 °C for 5 min. The sample was then subjected to electrophoresis through 10% denaturing polyacrylamide (19:1 acrylamide:bisacrylamide) with 7.5 M urea in 0.5× TBE running buffer. Electrophoresis was for 2 h at 19V/cm. The conjugated species was identified by gel shift via UV shadowing over a fluorescent TLC plate and excised from the gel with a cleaned razor blade. The oligonucleotide was eluted from the gel by overnight incubation in 400 µL 2× PK buffer (100 mM Tris-HCl, pH 7.5), 200 mM NaCl, 2 mM EDTA, 1% SDS) at 37 °C with rotation. The supernatant was then transferred and nucleic acids were extracted with 400 µL phenol:chloroform (1:1). The upper aqueous phase was transferred and precipitated with EtOH as described above.

**1.3 Conjugation of 5'-amino-modified oligodeoxyribonucleotide with tryptophan NHS ester**

Nα-Boc-l-tryptophan NHS ester (Trp-NHS) (Sigma-Aldrich, 15516) stock was prepared in anhydrous DMSO at 14 mM and air was evacuated with excess argon before sealing in air-tight tubes. Trp-NHS stock (25 µL) was combined with 50 µL NaB buffer (0.224 M $Na_2B_4O_7 \cdot 10H_2O$, pH 8.5), 30 µL 1 mM LJM-6543, and 20 µL water. The solution was mixed thoroughly by vortex mixing and incubated overnight at room temperature.

The reaction was then combined with 450 µL EtOH and 50 µL 3 M NaOAc and chilled on dry ice for 10 min. The sample was subjected to centrifugation at 17,000 × g for 20 min and supernatant was removed. The precipitated pellet was washed with 500 µL 70% EtOH and subjected to centrifugation again for 5 min. Supernatant was again removed and the pellet was air-dried at 37 °C. After drying, the conjugated oligo was resuspended in 100 µL water.

The precipitated oligonucleotide was combined with 100 µL formamide and heated at 90 °C for 5 min. The sample was then subjected to electrophoresis through 10% denaturing polyacrylamide (19:1 acrylamide:bisacrylamide) with 7.5 M urea in 0.5× TBE running buffer. Electrophoresis was for 2 h at 19 V/cm. The conjugated species was identified by gel shift via UV shadowing over a fluorescent TLC plate and excised from the gel with a cleaned razor blade. The oligonucleotide was eluted from the gel by overnight incubation in 400 µL 2× PK buffer (100 mM Tris-HCl, pH 7.5), 200 mM NaCl, 2 mM EDTA, 1% SDS) at 37 °C with rotation. The supernatant was then transferred and nucleic acids were extracted with 400 µL phenol:chloroform (1:1). The upper aqueous phase was transferred and precipitated with EtOH as described above.

**1.4 Mass Spectrometry**

**General Materials.** Nuclease P1 was purchased from US Biological Life Sciences and reconstituted according to the manufacturer's protocols. Calf intestinal alkaline phosphatase was purchased from Sigma. Spin filters (10 kDa molecular weight cut off) were purchased from VWR.

**Digestion of DNA oligonucleotides for LC-MS analysis.** Each oligonucleotide (100 pmol, 78 µL) was incubated with nuclease P1 (1.5 U, 3 µL) in 30 mM ammonium acetate pH 5.3 and 0.5 mM $ZnCl_2$ (90 µL total reaction volume) for 2 h at 55 °C. The reaction mixture was diluted with Tris-HCl (10 mM final concentration, pH 8.0, 9 µL) and incubated with calf intestinal alkaline phosphatase (51 U, 3 µL) for 2 h at 37 °C. Enzymes were removed by passing the mixture through a VWR 10 kDa spin filter with centrifugation at 12,000 g for 12 min at 4 °C). The solution was lyophilized to dryness and resuspended in $H_2O$ (50 µL).

**Orbitrap LC-MS of hydrolyzed oligonucleotides.** Hydrolyzed oligonucleotides (~20 pmol per 10 µL injection) were analyzed on a Dionex Ultimate 3000 UHPLC system equipped with a Synergi Fusion RP column (2.5 µm particle size, 100 Å pore size, 100 mm length, 2 mm inner diameter). The HPLC was coupled to a Thermo Fisher Q Exactive Hybrid Quadrupole-Orbitrap mass spectrometer. The column was eluted at 0.35 mL/min at 35 °C with a linear gradient of 3-65% acetonitrile in solvent A (5 mM ammonium acetate pH 5.3) over 35 min. The column was rinsed with 95% acetonitrile in solvent A for 1 min, and then the initial conditions were regenerated by rinsing the column with 97% solvent A for 3 min. High resolution mass spectra were obtained by hybrid quadrupole-Orbitrap mass spectrometry with the following parameters: sheath gas flow rate, 50 L/min; aux gas flow rate, 15 L/min; sweep gas flow rate, 3 L/min; spray voltage, 4.20 kV; and capillary temperature, 275 °C. Collision-induced dissociation was achieved using a collision energy of 40 V.

### 1.5 Bacterial Expression and Purification of TurboID Protein

The TurboID-His6 pET21a plasmid was transformed into T7 Express *lysY/I$^q$* Competent *E. coli* (New England BioLabs, C3013I). A colony was grown overnight in 5 mL LB. The entire culture was seeded into 500 mL LB and grown to an $OD_{600}$ value of 0.6. Protein expression was induced with 1 mM IPTG for 3 h. The bacteria were harvested by centrifugation at 6,000 × g for 15 min at 4˚C. Pelleted cells were resuspended in 40 mL lysis buffer (0.5 M NaCl, 10% glycerol, 20 mM HEPES, pH 8, 1 mM EDTA, 0.1% NP-40, 20 mM ß-mercaptoethanol, and 1 mM PMSF), lysed with a high-pressure microfluidizer Emulsiflex C-5 (Avestin Inc.), and subjected to centrifugation at 12,000 × g for 30 min at 4˚C. The supernatant was then passed through a Ni-NTA agarose column (Qiagen, 30230). The column was washed with 10 mL lysis buffer and with 150 mL wash buffer (20 mM HEPES, pH 8, 1 mM EDTA, 10% glycerol, 500 mM NaCl). A pre-elution was performed with 6 mL of 15 mM imidazole in wash buffer, and 2 mL fractions were collected. This was followed by an elution with 14 mL of 100 mM imidazole in wash buffer and 2 mL fractions were collected. Fraction samples were prepared with NuPAGE LDS Sample Buffer (4×) (Invitrogen, NP0007) and NuPAGE Sample Reducing Agent (10×) (Invitrogen, NP0004). These were then loaded onto a NuPAGE 10% Bis-Tris gel (Invitrogen, NP0303BOX), run in MES SDS buffer (Invitrogen, NP0002), and stained with Imperial Protein Stain (Thermo Scientific, 24615) following manufacturer's instructions. Pre-elution fractions 1-3 and elution fractions 1-3 were pooled, and the buffer was exchanged twice to remove imidazole with Vivaspin 20, 10,000 MWCO ultrafiltration unit (Sartorius, VS2002). The concentration of TurboID was determined using a Qubit 3 Fluorometer (Invitrogen, Q33216) and Qubit Protein Assay Kit (Invitrogen, Q33211).

### 1.6 TurboID biotinylation activity assay by Western blot

Radical biotinylation reactions were performed with and without BT as controls. Reactions (40 µL) contained 10 mM potassium phosphate, pH 6, 12.5 µM HRP (Thermo Scientific, 31490), freshly prepared 500 µM $H_2O_2$ (Sigma, 216763), and 100 µM biotin tyramide (ApexBio, A8011) when applicable. Reactions were incubated at 45 ºC for 1.5 h with agitation at 1000 rpm (Benchmark Scientific, model H5000-HC).

Biotin ligase reactions (20 µL) contained HEPES buffer (20 mM HEPES, pH 8, 500 mM NaCl), 5 mM $MgCl_2$, 5 µM TurboID, 10 mM ATP (Thermo Scientific, R0441), and 50 µM biotin (Sigma, B4639). Reactions were incubated at 37˚C for 1 h.

Samples were prepared with NuPAGE LDS Sample Buffer (4×) (Invitrogen, NP0007) and NuPAGE Sample Reducing Agent (10X) (Invitrogen, NP0004). These were then loaded onto a NuPAGE 10% Bis-Tris gel (Invitrogen, NP0303BOX) and run in 1× MES SDS buffer (Invitrogen, NP0002) at 150 V for 1 h. The gel was blotted onto a PVDF membrane (Bio-Rad, 1620174) using an XCell II Blot Module (Invitrogen, EI9051) in 1× NuPAGE transfer buffer (Invitrogen, NP0006) containing 20% methanol. Blotted membrane was briefly washed in TBST (50 mM Tris-HCl, pH 7.4, 150 mM NaCl, 0.1% Tween-20) and treated with blocking buffer (5% dry milk and 1% BSA in TBST) overnight at 4˚C with gentle rocking. The membrane was incubated with Streptavidin Alexa Fluor 647 (Invitrogen, S21374) diluted 1:500 in long term storage buffer (4% BSA and 0.2% NaAzide in TBST) for 2 h at room temperature with gentle rocking. The membrane was washed with TBST three times for 20 min each at room temperature. Membranes were analyzed with an Amersham Typhoon laser-scanner platform (Cytiva) using the Alexa Fluor 647 channel.

**1.7 TurboID biotinylation assay**

Biotin ligase reactions (20 µL) contained 2.5 µM indicated oligonucleotide, HEPES buffer (20 mM HEPES, pH 8, 500 mM NaCl), 5 mM $MgCl_2$, 5 µM TurboID, 10 mM ATP (Thermo Scientific, R0441), and 50 µM biotin (Sigma, B4639). When applicable, glycine was included as a competitor at 12.5 mM and water was subtracted to maintain a constant final reaction volume. Reactions were incubated at 37 ºC for 24 h. Reactions were then diluted by addition of 380 µL water and extracted with 400 µL phenol:chloroform:isoamyl alcohol (25:24:1) (Invitrogen, 15593031). Aqueous phases were combined with 0.1 volume 3 M sodium acetate, 1 µl glycogen (Thermo Scientific, R0561) and 2.5 volumes ice-cold 100% ethanol. This mixture was chilled on dry ice for 20 min and subjected to centrifugation for 20 min at 17,000 × g. Samples were washed with 400 µL ice-cold 70% ethanol followed by centrifugation for 10 min at 17,000 × g and air drying. Pellets were resuspended in water (15 µL), and either streptavidin (5 µL of 182 µM stock; Invitrogen, 434301) or water (5 µL) was added with incubation for 1.5 h at 37 ºC with agitation at 1000 rpm (Benchmark Scientific, H5000-HC). Samples were then analyzed by electrophoresis. When applicable, streptavidin was incubated with 5 µL of 1 mM biotin prior to adding to resuspended DNA pellet, which was resuspended in 10 µL water to maintain a constant final volume. For analysis under native conditions, samples were supplemented with 0.2 volume loading buffer and subjected to electrophoresis at 22.5 V/cm for 35 min through native 10% polyacrylamide (19:1 acrylamide:bisacrylamide) gels. Gels were imaged on a Typhoon fluorometric imager using the FAM channel.

**2.0 Supplemental data**

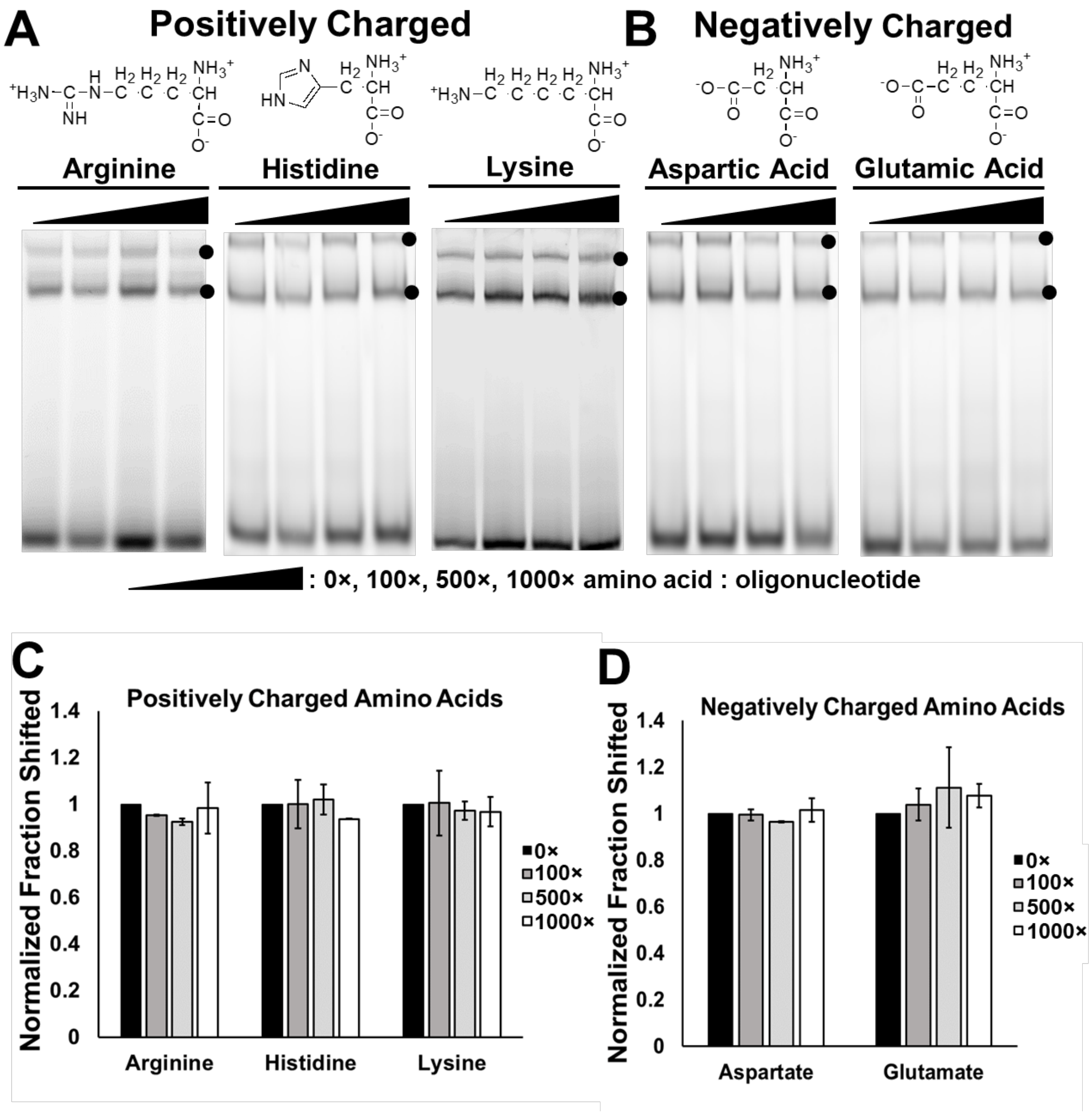


**Figure S1. Positively and negatively charged amino acids do not react with phenoxy radical BT intermediates.** A) Excess Arg, His, or Lys are added to in vitro HRP-mediated biotinylation reaction and allowed to compete for radical reactivity. Covalently biotinylated oligonucleotides are detected by streptavidin gel shift, which is quenched in the presence of reactive competitors. B) Excess Asp and Glu are added to in vitro HRP-mediated biotinylation reaction and allowed to compete for radical reactivity. C) Quantification of gel shifted material observed in panel A. D) Quantification of gel shifts observed in panel B.

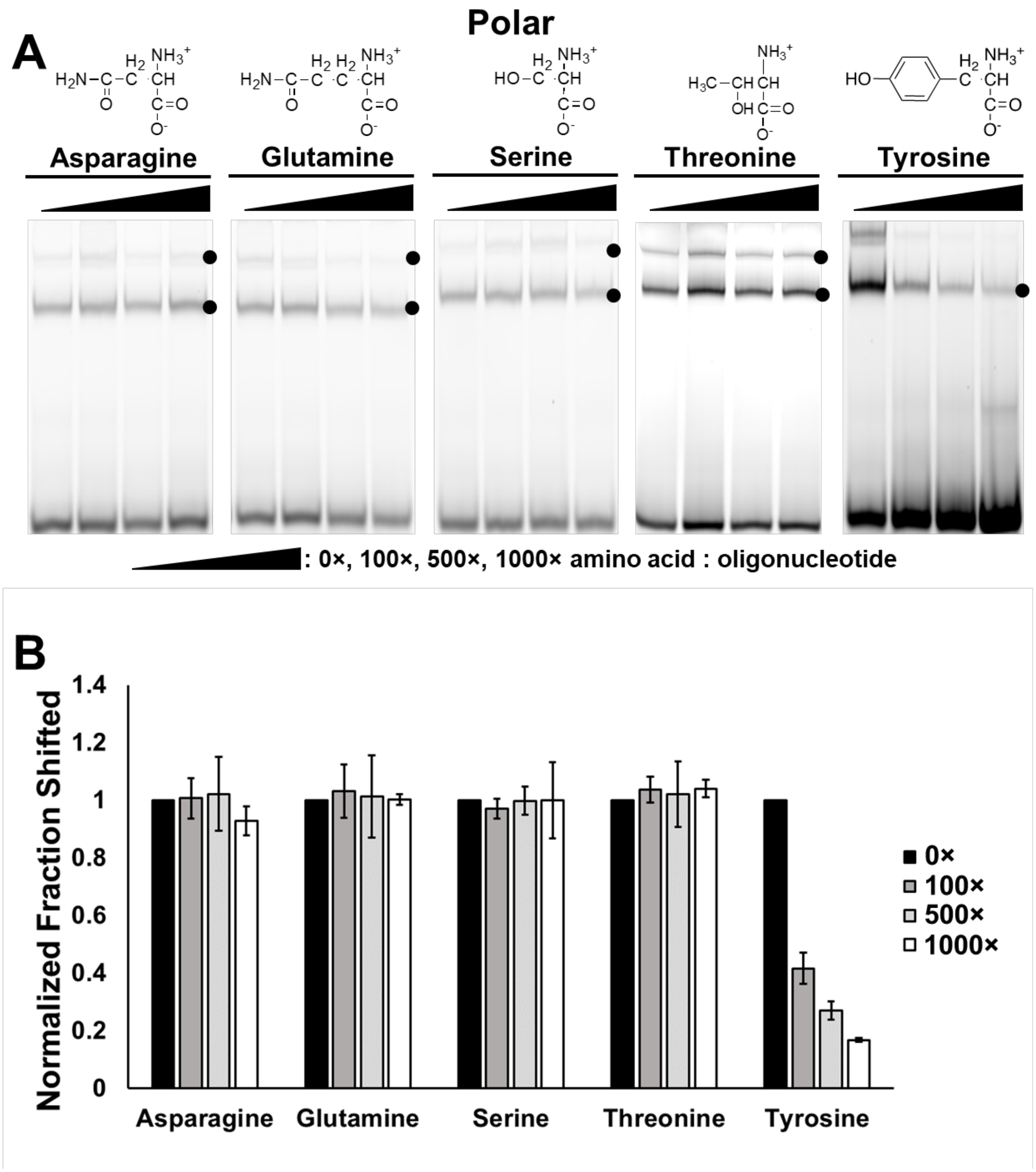


**Figure S2. Tyrosine competes for phenoxy radical biotinylation.** A) Excess Asn, Gln, Ser, Thr, and Tyr are added to in vitro HRP-mediated biotinylation reaction and allowed to compete for radical reactivity. Covalently biotinylated oligonucleotides are detected by streptavidin gel shift, which is quenched in the presence of reactive competitors. B) Quantification of gel shifts observed in panel A.

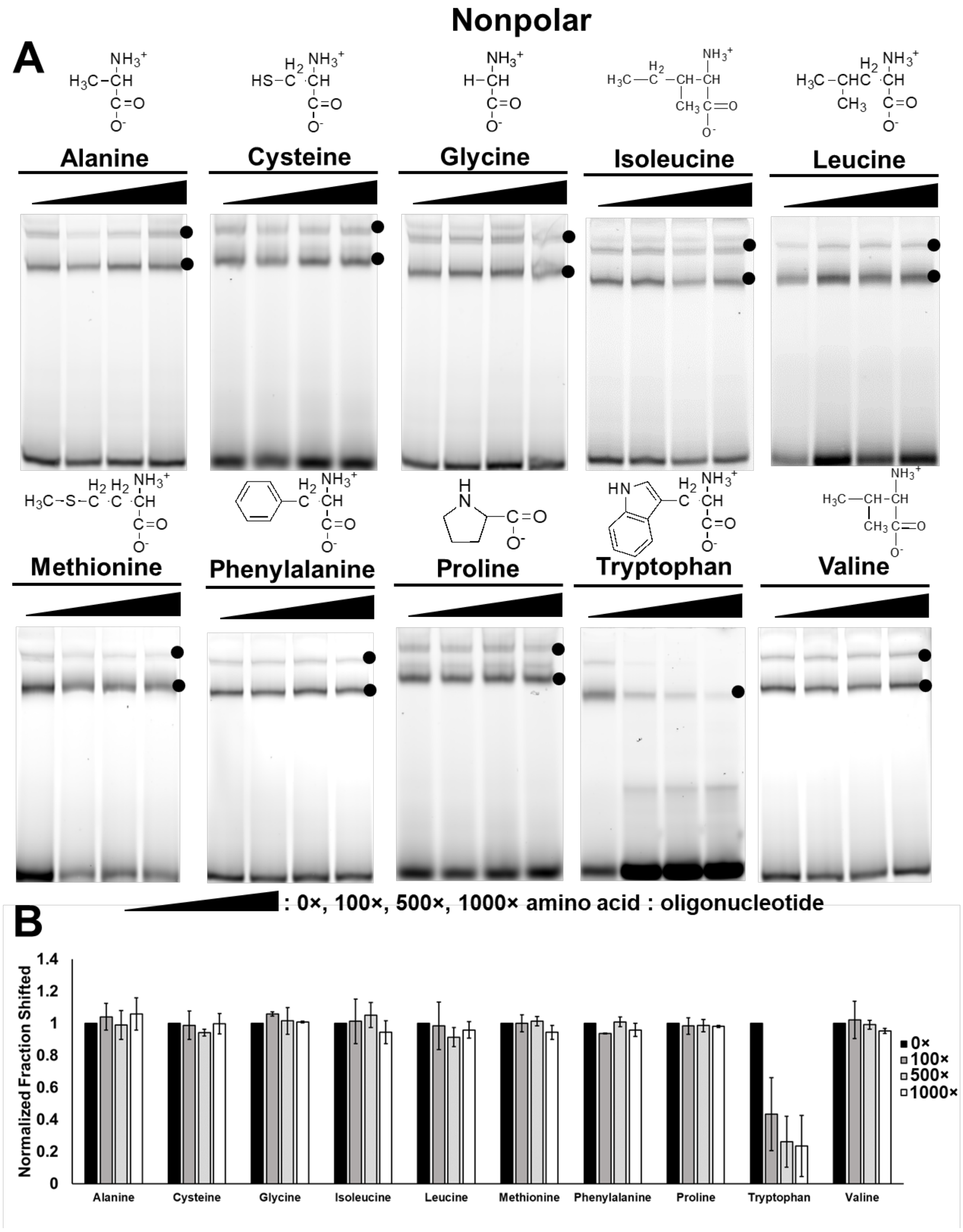


**Figure S3. Tryptophan competes for phenoxy radical biotinylation.** A) Excess nonpolar amino acids are added to *in vitro* HRP-mediated biotinylation reaction and allowed to compete for radical reactivity. Covalently biotinylated oligonucleotides are detected by streptavidin gel shift, which is quenched in the presence of reactive competitors. B) Quantification of gel shifts material observed in panel A.

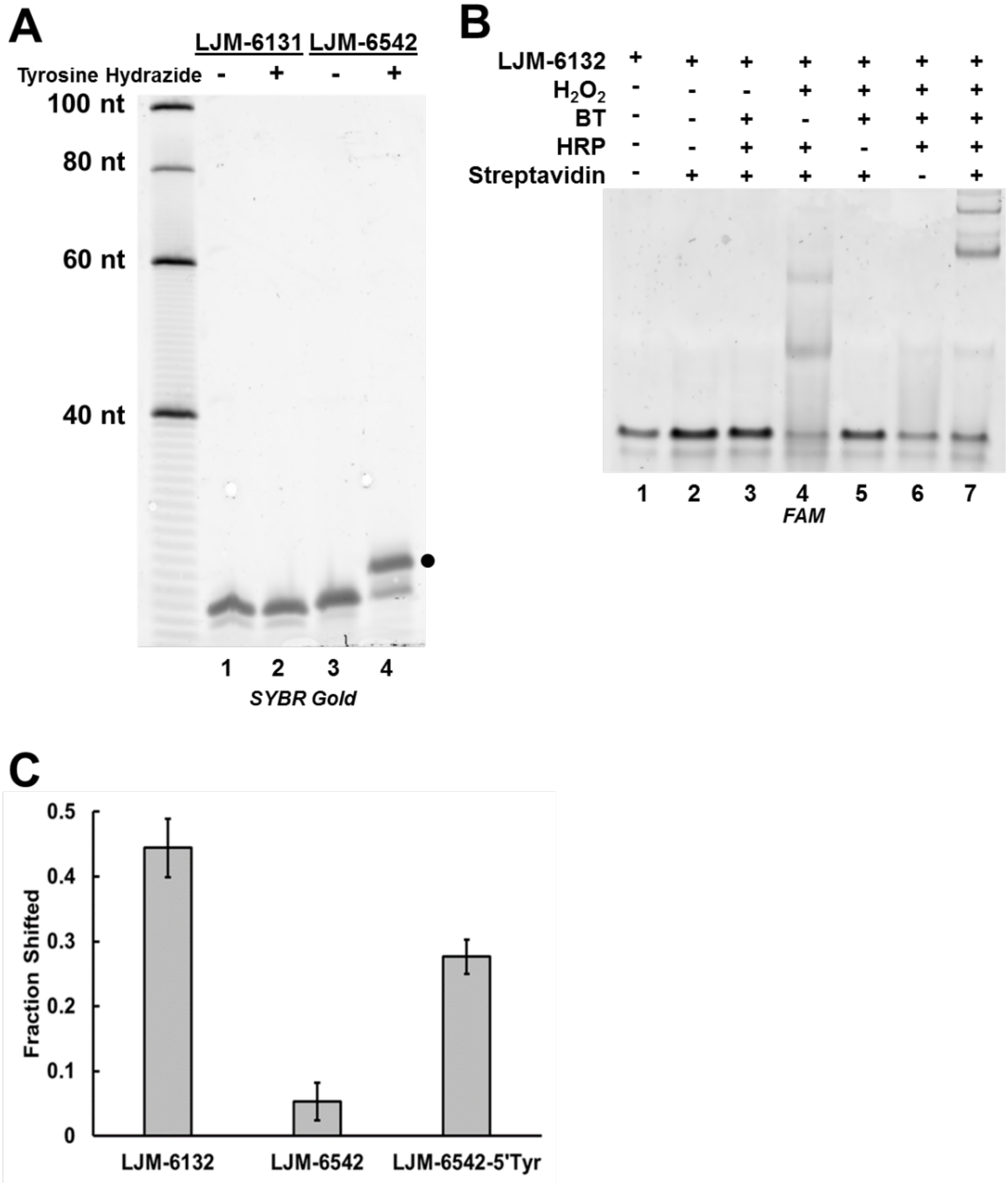


**Figure S4. Conjugation of tyrosine hydrazide to LJM-6542 and in vitro biotinylation of LJM-6132**. A) Unmodified oligonucleotide LJM-6131 is not conjugated or gel shifted in the presence of tyrosine hydrazide. LJM-6542, a 5' aromatic aldehyde-modified oligonucleotide, reacts with tyrosine hydrazide to covalently attach tyrosine and is gel shifted as a result (black dot). B) 5' fluorescein modified oligonucleotide LJM-6132 is biotinylated and streptavidin gel shifted only in the presence of $H_2O_2$, BT, and HRP (lane 7). BT-free gel shifts are observed due to the formation of fluorescein-fluorescein conjugates (lane 4). C) Quantification of LJM-6132, LJM-6542, and LJM-6542-5'Tyr streptavidin gel shifts.

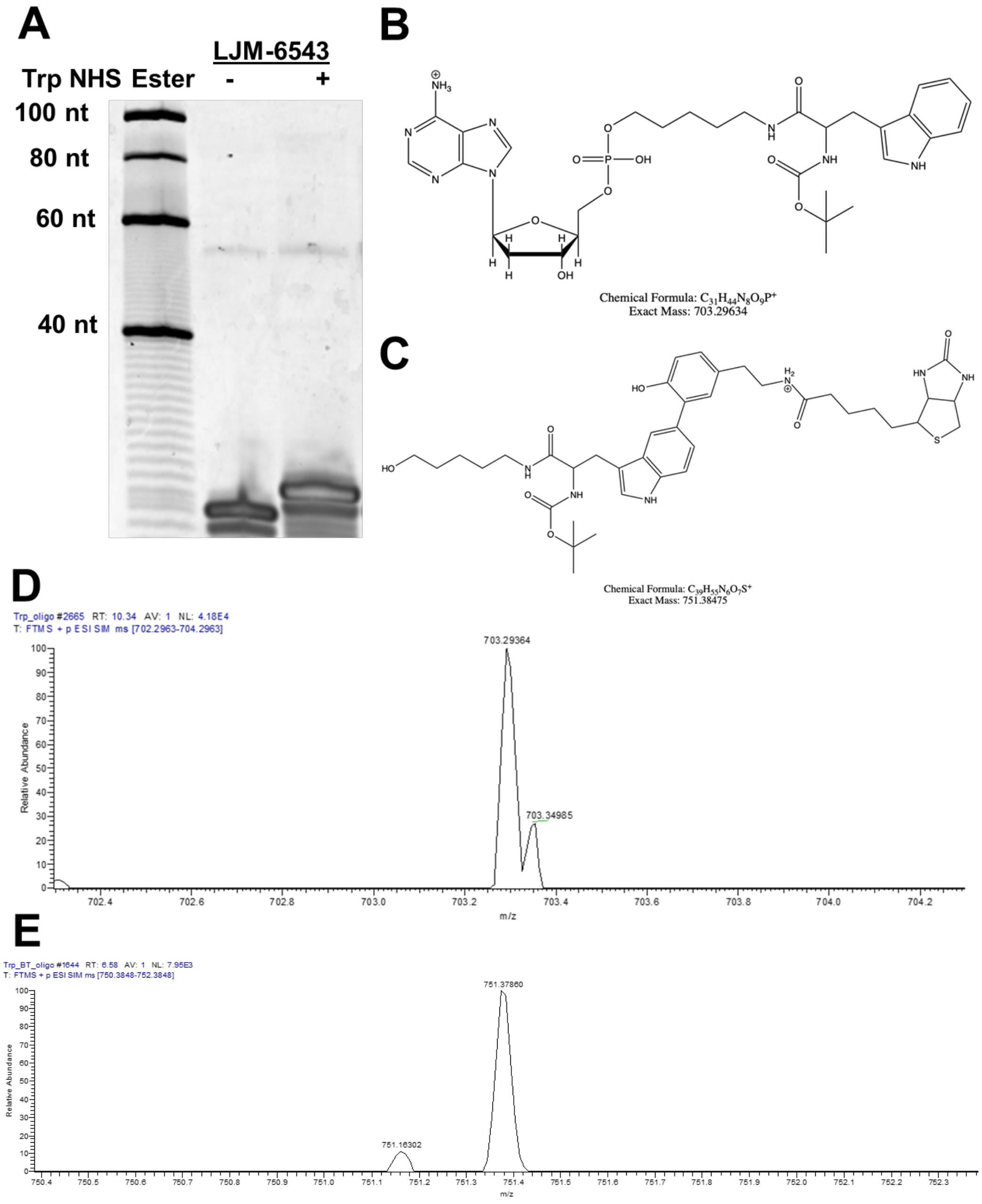


**Figure S5. Conjugation of Trp-NHS ester to LJM-6543 and analysis of products of HRP-free biotinylation.** A) 5' primary amine-modified oligonucleotide LJM-6543 is covalently conjugated to tryptophan-NHS ester as observed by gel shift in denaturing polyacrylamide. B) Predicted product of tryptophan-NHS ester conjugation to LJM-6543. C) Predicted structure of biotinylated products resulting from HRP-free biotinylation of LJM-6543-5'Trp. D) Mass spectra of the predicted product in panel B confirms that conjugation results in the expected product mass. E) Mass spectra of the predicted product in panel C confirms that HRP-free biotinylation results in the expected product mass.

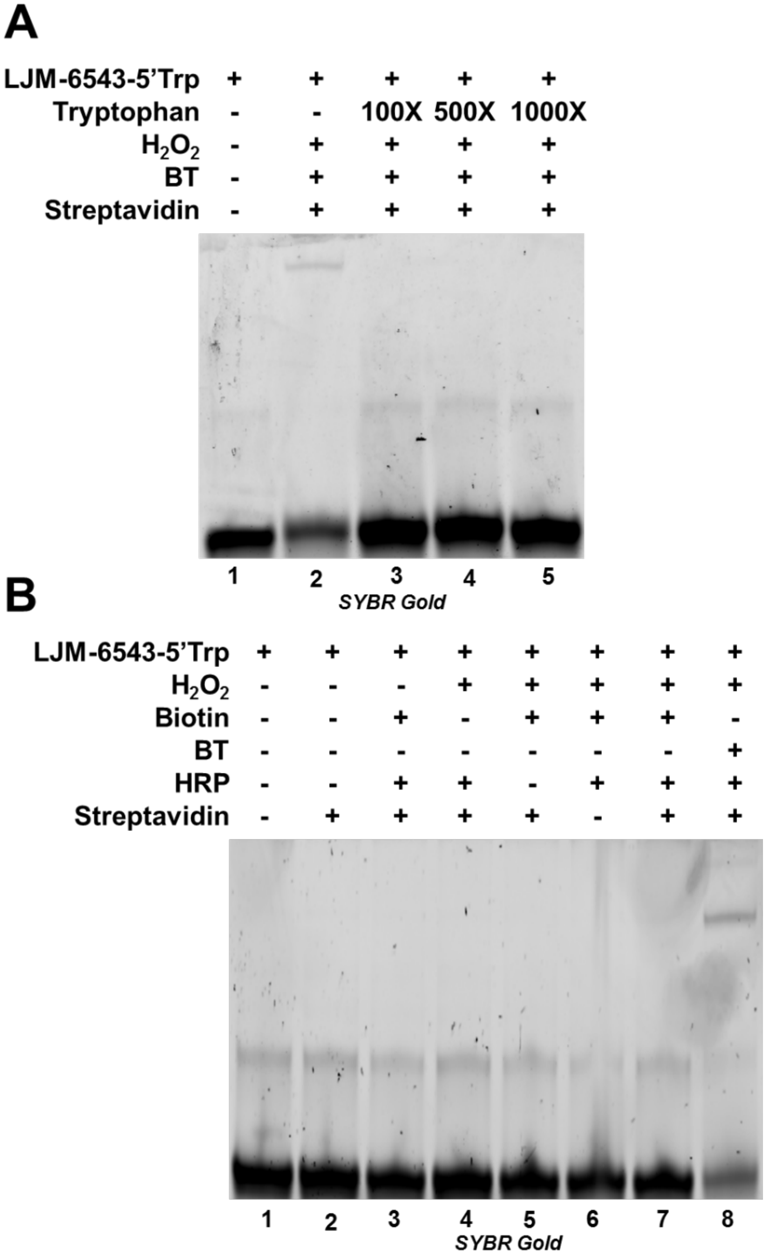


**Figure S6. Characterization of HRP-free biotinylation reactions of LJM-6543-5'-Trp.** A) Tryptophan is added to HRP-free biotinylation reactions of LJM-6543-5'Trp in up to 1000× molar excess of oligonucleotide. Excess tryptophan quenches the gel shift observed in lane 2, demonstrating that the Boc modification included in Trp-NHS ester does not participate in the reaction. B) LJM-6543-5' Trp can only be biotinylated by biotin tyramide, not biotin, in standard or HRP-free reaction conditions.

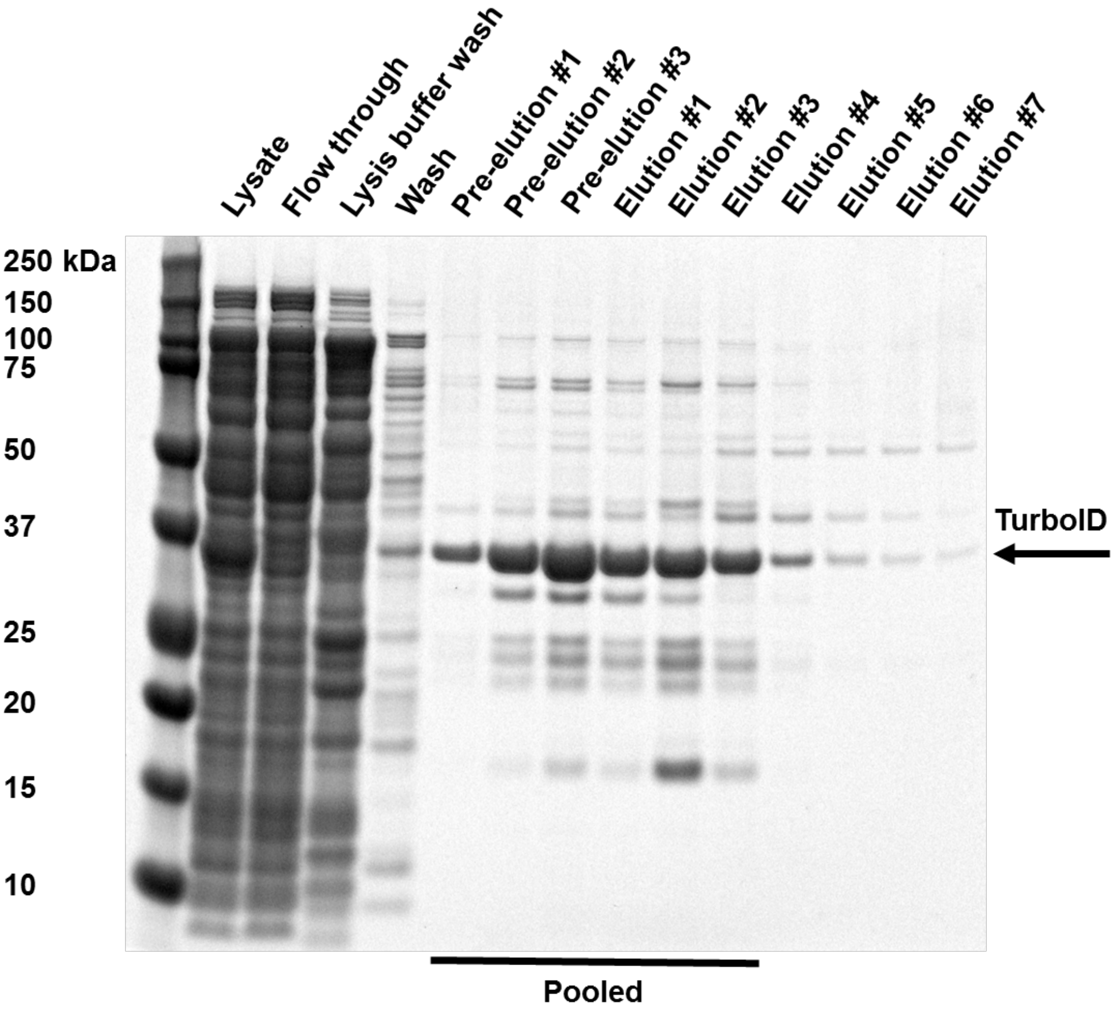


**Figure S7. Purification of TurboID.** Coomassie staining of protein purification fractions during Ni-NTA agarose column purification of TurboID. Pooled fractions are those combined for use in this study.

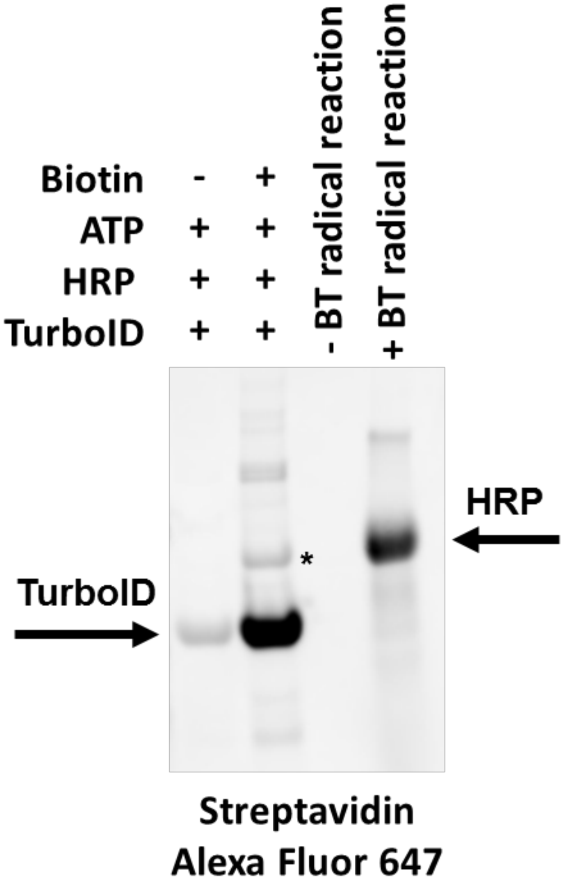


**Figure S8. Validation of TurboID biotin ligase activity by streptavidin Western blot.** Low levels of TurboID self-biotinylation are detectable in purified protein without the addition of biotin to *in vitro* reactions. This is an expected result of basal *in vivo* self-biotinylation *E. coli* before purification; addition of biotin to *in vitro* reactions greatly increases intramolecular biotinylation. Biotinylation of supplemented HRP, indicated with an asterisk, shows intermolecular biotin ligase activity. Other contaminating proteins were also biotinylated. Previously described self-biotinylation of HRP by biotin tyramide is seen only in the presence of biotin tyramide and $H_2O_2$.

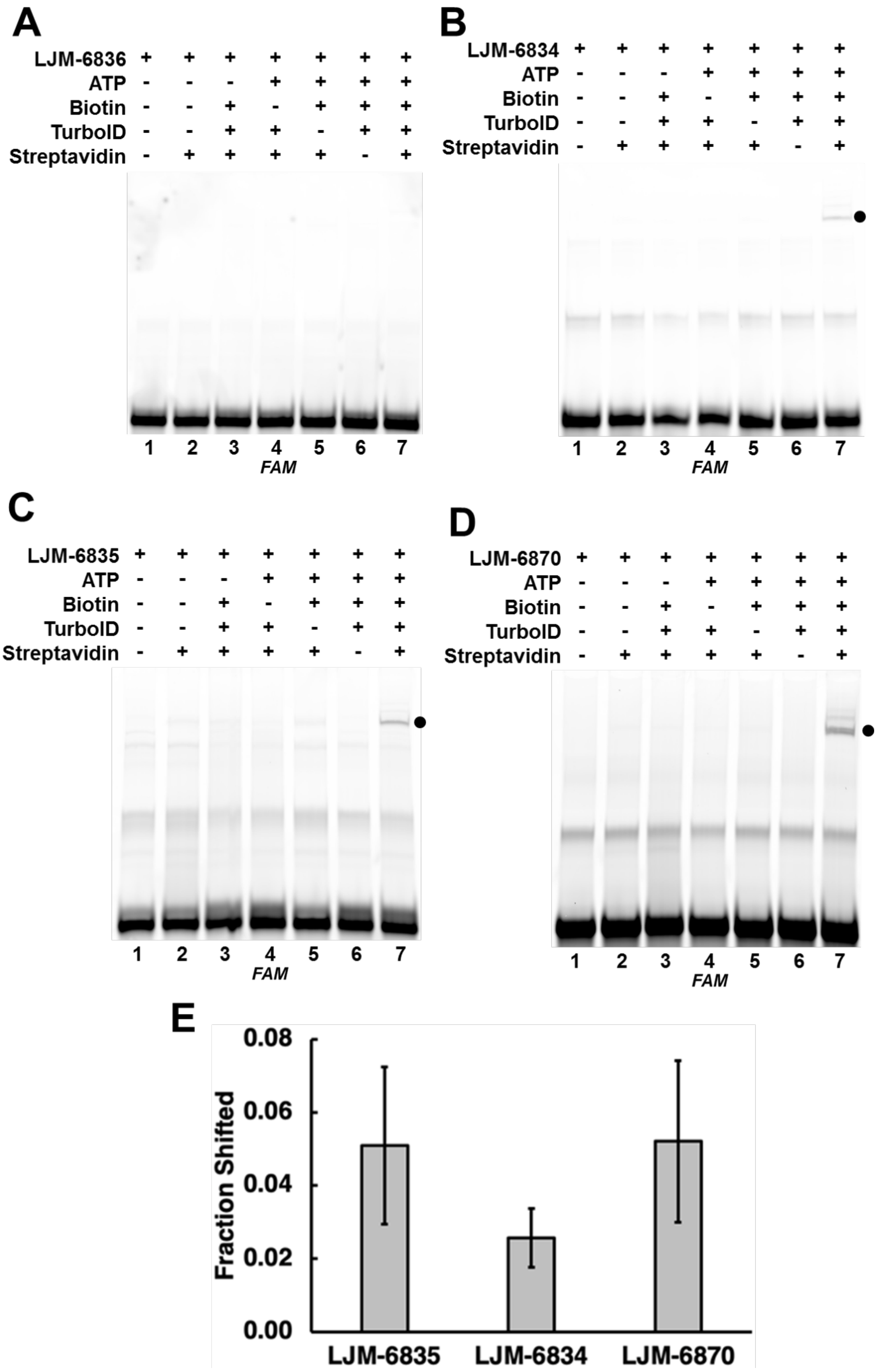

A
LJM-6836
ATP
Biotin
TurboID
Streptavidin
1 2 3 4 5 6 7
FAM
B
LJM-6834
ATP
Biotin
TurboID
Streptavidin
1 2 3 4 5 6 7
FAM
C
LJM-6835
ATP
Biotin
TurboID
Streptavidin
1 2 3 4 5 6 7
FAM
D
LJM-6870
ATP
Biotin
TurboID
Streptavidin
1 2 3 4 5 6 7
FAM
E
Fraction Shifted
0.08
0.06
0.04
0.02
0.00
LJM-6835
LJM-6834
LJM-6870

**Figure S9. Unmodified oligonucleotides are not biotinylated by TurboID and all surveyed amino acid modifications react similarly.** A) An oligonucleotide lacking any amino groups (LJM-6836) is not detectably biotinylated in the presence ATP, Biotin, and TurboID. B) Biotinylation of 5’ primary amine modified oligonucleotide (with six-carbon spacer) LJM-6834 (black dot) occurs only in the presence of ATP, Biotin, and TurboID. C) Biotinylation of 5’ primary amine modified oligonucleotide (with twelve-carbon spacer) LJM-6835 (black dot) occurs only in the presence of ATP, Biotin, and TurboID. D) Biotinylation of oligonucleotide containing 5’ primary amine modification with six-carbon spacer and two internal primary amine modified deoxythymidine nucleotides LJM-6870 (black dot) occurs only in the presence of ATP, Biotin, and TurboID. E) Quantification of observed gel shifts.

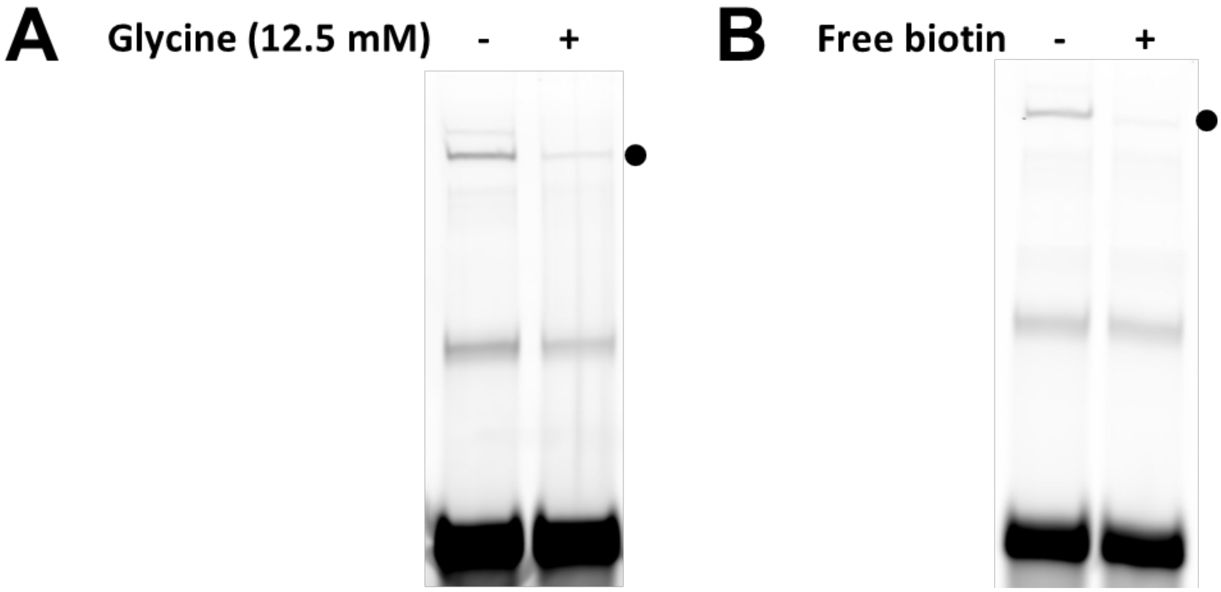


**Figure S10. Biotinylation reaction catalyzed by TurboID can be quenched with excess glycine, and the biotinylation specificity of gel shift is confirmed by free biotin competition.** A) Biotinylation and streptavidin gel shift of LJM-6834 is diminish by excess glycine included during *in vitro* TurboID biotinylation reaction mixtures as a primary amine competitor (black dot). B) When streptavidin is blocked with excess biotin prior to incubation with biotinylated LJM-6834, the gel shift is diminished (black dot).

**Supplemental Table S1.**

| **Serial number** | **Oligonucleotide sequence**[1] |
|---|---|
| LJM-6132 | /56-FAM/AGGGCTAAGGTCTGACTGGGAATGGGA |
| LJM-6542 | /5AhMC2/AGGGCTAAGGTCTGACTGGGAATGGGA |
| LJM-6543 | /5AmMC6/AGGGCTAAGGTCTGACTGGGAATGGGA |
| LJM-6834 | /5AmMC6/AGGGCTAAGGTCTGACTGGGAATGGGA/36-FAM/ |
| LJM-6835 | /5AmMC12/AGGGCTAAGGTCTGACTGGGAATGGGA/36-FAM/ |
| LJM-6836 | AGGGCTAAGGTCTGACTGGGAATGGGA/36-FAM/ |
| LJM-6870 | /5AmMC6/AGGGC/iAmMC6T/AAGGTCTGACTGGGAA/iAmMC6T/GGGA/36-FAM/ |
| LJM-6892 | /56-FAM/AGGGCTAAGGTCTGACTGGGAATGGGA/3ThioMC3-D/ |

[1]Modification abbreviations: /56-FAM/ is an isomer derivative of fluorescein with a six-carbon spacer at the 5' terminus end of the oligonucleotide. /5AmMC6/ is a primary amino group with a six-carbon spacer attached at the 5' terminus of the oligonucleotide. /5AhMC2/ is a *para*-aromatic aldehyde ether linked to a two-carbon spacer attached at the 5' terminus of the oligonucleotide. /36-FAM/ is an isomer derivative of fluorescein with a six-carbon spacer attached to the 3' terminus of the oligonucleotide. /5AmMC12/ is a primary amino group with a twelve-carbon spacer attached to the 5' end of the oligonucleotide. /iAmMC6T/is a deoxythymidine analog with a primary amine at the 5 position of the base. /56-FAM/ is an isomer derivative of fluorescein with a six-carbon spacer attached at the 5' terminus of the oligonucleotide. /3ThioMC3-D/ is a thiol with a three-carbon spacer attached at the 3' terminus of the oligonucleotide that is purchased in the oxidized form. All oligonucleotides with modifications were purchased from Integrated DNA Technologies.
